# Low frequency optogenetic deep brain stimulation of nucleus accumbens dopamine D1 or D2 receptor-containing neurons attenuates cocaine seeking selectively in male rats in part by reversing synaptic plasticity deficits

**DOI:** 10.1101/2023.01.23.524956

**Authors:** Sarah E. Swinford-Jackson, Matthew T. Rich, Phillip J. Huffman, Melissa C. Knouse, Arthur S. Thomas, Sharvari Mankame, Samantha J. Worobey, R. Christopher Pierce

## Abstract

**Background:** Clinically, deep brain stimulation (DBS) utilizes relatively high frequencies (>100 Hz). In preclinical models, 160 Hz stimulation of the nucleus accumbens in rodents prevents relapse of drug seeking. However, the ability of varied frequencies of accumbens DBS to attenuate drug seeking, and the neuronal subtype specificity of this effect, is unclear.

**Methods:** The present study examined the effect of DBS in the nucleus accumbens on neuronal plasticity and cocaine-primed reinstatement of cocaine seeking behavior in rats.

**Results:** Electrical DBS of the accumbens shell attenuated cocaine primed reinstatement across a range of frequencies in male rats, including as low as 12 Hz. The majority of nucleus accumbens neurons are medium spiny neurons (MSNs), which can be differentiated in terms of projections and effects on cocaine-related behaviors by expression of dopamine D1 receptors (D1DRs) or D2DRs. In slice electrophysiology experiments, 12 Hz electrical stimulation evoked long term potentiation (LTP) in eYFP labeled D1DR-MSNs and D2DR-MSNs from cocaine naive male and female rats. However, in rats that self-administered cocaine and underwent extinction training, a paradigm identical to our reinstatement experiments, electrical DBS only elicited LTP in D2DR-MSNs from male rats; this effect was replicated by optical stimulation in rats expressing Cre-dependent ChR2 in D2DR-MSNs. Low-frequency optogenetic-DBS in D1DR-containing or D2DR-containing neurons attenuated cocaine-primed reinstatement of cocaine seeking in male but not female rats.

**Conclusions:** These results suggest that administering DBS in the nucleus accumbens shell at lower frequencies effectively, but sex-specifically, suppresses cocaine craving, perhaps in part by reversing synaptic plasticity deficits selectively in D2DR-MSNs.

## Introduction

High rates of relapse remain a significant impediment to recovery in individuals with cocaine use disorder (1). Although effective pharmacological therapies are unavailable, DBS has been established as a treatment possibility for severe substance use disorders (2). First approved for Parkinson’s Disease, clinical studies using DBS now include other neurological and neuropsychiatric disorders including substance use disorders (2-5). Early clinical and preclinical studies on Parkinson’s Disease informed commonly used stimulation parameters, typically characterized by higher stimulation frequencies (i.e. greater than 100 Hz) (6, 7). Recent studies have begun to explore the therapeutic benefit of lower frequency DBS (8-11), although no clinical studies have yet applied these parameters to substance use disorder treatment.

Facets of cocaine use disorder can be modeled in rodents using drug self-administration paradigms. In rats, bilateral high frequency DBS of the nucleus accumbens shell attenuated cocaine primin1g- and cue-induced reinstatement of cocaine seeking (12-14), two common models of relapse. In contrast, in rats which expressed escalated cocaine self-administration, accumbens DBS actually increased cocaine taking (15). Preclinical DBS studies (12-14, 16) have typically employed the high frequency parameters most often used clinically (2, 17-22). However, low frequency accumbens DBS has also been shown to alter drug-related behaviors. For example, chronic unilateral nucleus accumbens shell DBS delivered at 20 Hz during abstinence from cocaine self-administration attenuated cocaine seeking (23).

The vast majority of accumbens neurons are medium spiny efferent neurons. These medium spiny neurons (MSNs) are all GABAergic but can be divided into two similar sized populations based on the expression of either D1DRs or D2DRs (for evidence of limited co-expression see (24)). Chemogenetic and optogenetic approaches have defined bidirectional roles for accumbens D1DR-MSNs and D2DR-MSNs such that activation of D1DR-MSNs potentiates various cocaine-mediated behaviors, whereas they are inhibited by stimulation of D2DR-MSNs (25-27). Recent studies have begun to explore the cell subtype-specificity of DBS to modulate behavior using optogenetic approaches. Mimicking high-frequency electrical DBS with optogenetic stimulation within nucleus accumbens neuronal subtypes indicated that 130 Hz opto-DBS of D2DR-containing, but not D1DR-containing neurons attenuated cocaine seeking in male rats (28). Low frequency opto-DBS (12 Hz) of neurons projecting from the prefrontal cortex onto accumbens D1DR-MSNs blocked cocaine-evoked locomotor sensitization and altered long-term synaptic plasticity; importantly, these effects were only elicited by 12 Hz electrical DBS when administered in combination with a D1DR antagonist (29). These studies suggest that MSN subtype-specific manipulation using opto-DBS may be used to disentangle the intricate and even opposing effects of electrical DBS which impinge on cocaine-related behaviors.

The present study aimed to investigate the effect of low frequency accumbens shell DBS on cocaine seeking and to elucidate the MSN subtype-specific mechanisms underlying the suppression of cocaine-primed reinstatement of cocaine seeking. Nucleus accumbens DBS attenuated cocaine seeking over a range of frequencies in male rats, including frequencies as low as 12 Hz. To parse out potential mechanisms underlying this effect, we focused on changes in long term synaptic plasticity in accumbens MSNs from naive and cocaine-trained, male and female, D1DR-Cre and D2DR-Cre rats (28, 30-32). Slice electrophysiology experiments showed that low frequency stimulation elicited long term potentiation (LTP) in both accumbens MSN subtypes from naive male and female rats. However, in cocaine-experienced rats, 12 Hz electrical or optical stimulation selectively evoked LTP only in D2DR-MSNs of males.

Surprisingly, this effect was sex-specific in that 12 Hz opto-DBS of the accumbens shell attenuated cocaine-primed reinstatement in male, but not female rats.

## Materials and Methods

### Animals and housing

Male Sprague Dawley rats (250-300g) were procured from Taconic (Germantown, NY). LE-Tg(Drd1a-iCre)3Ottc (RRRC#:00767; D1DR-Cre) and LE-Tg(Drd2-iCre)1Ottc (RRRC#:00768; D2DR-Cre) male founders were generated by the NIDA Optogenetics and Transgenic Technology Core (now Transgenic Rat Project, NIDA IRP, Bethesda, MA) and obtained from the Rat Resource and Research Center (Colombia, MS). Breeders were backcrossed frequently with female Long Evans (Charles River Laboratories, Wilmington, MA) to prevent genetic drift. Adult (PND 60-240) male and female D1DR-Cre or D2DR-Cre rats bred in house were used for experiments. Rats were individually housed with food and water available *ad libitum*. A 12/12 hr light/dark cycle was used with the lights on at 6:00 a.m. All experimental procedures were performed during the light cycle. All experimental procedures were consistent with the ethical guidelines of the US National Institutes of Health and were approved by the Institutional Animal Care and Use Committee.

### Surgery

An indwelling silastic catheter (SAI Infusion Technologies, Libertyville, Il) was inserted into the jugular vein and sutured in place as previously described (33, 34). Following catheter insertion, rats received intra-nucleus accumbens shell implantation of electrodes (12, 13) or viral vector infusion with fiber optic implantation (28), as previously described.

### Ex vivo electrophysiology

Rats expressing Cre-dependent eYFP or ChR2 in the nucleus accumbens were collected for whole-cell patch-clamp recordings (28, 35-37) using methods designed to improve neuronal health in adult rodents (38, 39). MSNs were identified by their morphology and low resting membrane potential (−70 to −85 mV), and D1DR- or D2DR-containing neurons were further identified by eYFP expression. All recordings were performed in whole-cell voltage-clamp or current-clamp mode using a MultiClamp 700B amplifier and Digidata 1440A Digitizer (Molecular Devices, San Jose, CA). A concentric bipolar stimulating electrode (FHC, Bowdoin, ME) was placed over axon fibers in close proximity to the recorded neuron. These projections were stimulated using 0.1 ms pulses (at 0.05 Hz) delivered from an isolated current stimulator (A-M instruments; Digitimer Ltd, Hertfordshire, England), and the evoked excitatory postsynaptic currents (EPSCs) were recorded. For experiments involving optical stimulation, ChR2+ neurons were directly stimulated using blue light (473 nm) from either a DPSS laser (OptoEngine, Midvale, UT) or an external 4 channel LED driver (Thor Labs, Newton, NJ). The laser/LED intensity was adjusted to the minimal amplitude (∼1 mW or 10-50 mA) necessary to elicit action potential firing with a single 5 ms pulse. Following collection of baseline EPSCs, neurons received 10 min of electrical or optical 12 Hz stimulation. A stimulation duration of 10 minutes corresponds with peak responding in a cocaine-primed reinstatement session and preserves the health of patched neurons. EPSCs were continuously recorded at 0.05 Hz for ≥ 40 min. The baseline amplitudes were compared to the EPSC amplitude following the DBS stimulation protocol (averaged over 10 mins pre-stim and last 10 min post-stim). Pair-pulsed ratios of evoked EPSCs and spontaneous activity before and after the DBS stimulation protocol were measured to gain insight into alterations in presynaptic function. All analyses of intracellular recordings were performed with Clampfit 10 (Molecular Devices). The experimenter was blind to treatment conditions during all electrophysiological recordings and analyses. For all experiments, series resistance was 10–25 MΩ, uncompensated, and monitored continuously during recording. Cells with a change in series resistance beyond 20% were excluded. Synaptic currents were filtered at 3 kHz, amplified 5 times, and then digitized at 20 kHz.

### Cocaine self-administration, extinction, and reinstatement

Rats were placed in operant conditioning chambers (Med-Associates, East Fairfield, VT) and allowed to press the active lever for intravenous cocaine infusions (0.25 mg of cocaine in 59 µL of saline, infused over 5 seconds, maximum of 30 infusions/session); inactive lever responses produced no scheduled consequences. Rats began training on a fixed ratio 1 (FR1) schedule of reinforcement. When stable responding was achieved on the FR1 schedule (less than 15% variation over three consecutive days), rats were switched to an FR5 schedule. A 20 s timeout period during which active lever responses had no scheduled consequences followed each cocaine infusion. After 21 days of cocaine self-administration sessions, rats underwent an extinction phase during which cocaine was replaced with 0.9% saline. Daily 2-hour extinction sessions were conducted until active lever responding was <20% of the responses averaged over the last three days of cocaine self-administration. Rats designated for *ex vivo* slice electrophysiology were evaluated at this timepoint to recapitulate the neuronal status of rats immediately prior to a reinstatement session. Cocaine seeking was reinstated by non-contingent administration of cocaine (10 mg/kg, i.p.) immediately prior to the initiation of the reinstatement session, during which satisfaction of the response requirement is not reinforced. Each reinstatement test day was followed by extinction sessions until responding was again <20% of the responses achieved during self-administration across two consecutive sessions.

### Deep brain stimulation

Each rat underwent multiple cocaine-primed reinstatement sessions during which intra-accumbens DBS was delivered at 150 μA and 12 Hz, 60Hz, 100 Hz, 130 Hz, or 0 μA (sham) in a within-subjects counterbalanced order to obviate any effect of test order.

Alternating current with biphasic symmetrical pulses (60 µs pulse width) and a 150 µA stimulation intensity were used for all DBS frequencies. DBS was initiated concurrently with the onset of the reinstatement session and administered bilaterally and continuously throughout the 2-hour reinstatement session, as described previously (12-14, 16).

### Opto-DBS

Each rat underwent two reinstatement sessions, during which 473 nm light stimulation delivered at 12 Hz with a 5ms pulse width or sham opto-DBS (patch cables attached but 0 mW delivered) was administered in a within-subjects counterbalanced fashion. Opto-DBS was administered continuously during the 1-hour reinstatement sessions, as described previously (28). Light from a 473 nm laser (OptoEngine, Midvale, UT) was split by a rotary joint (Doric Lenses, Quebec, Canada) delivered bilaterally through 200 µm fiber optic patch cables (Thor Labs) connected to the implanted ferrules. Frequency was modulated by a Master 8 pulse generator (AMPI, Jerusalem, Israel) and laser output was calibrated to 1 mW of light in the accumbens to prevent any effect of heat.

### Statistics

Statistical analysis was performed in Prism 9.0 with alpha set at p<0.05. Rats that maintained catheter patency throughout the duration of cocaine self-administration were included in behavioral analyses. Only rats with verified nucleus accumbens shell placement for AAV infusions alone or with fiber optic or electrodes were included in behavioral or electrophysiological experiments. Statistical tests are described within Figure Legends.

## Results

### Nucleus accumbens DBS attenuates cocaine seeking across a range of frequencies

Male Sprague Dawley rats were allowed to self-administer cocaine for 21 days on an FR1-5 schedule, as described previously (12, 13, 16, 28). Following cocaine self-administration training and extinction of lever responding, cocaine seeking was assessed in cocaine-primed reinstatement sessions during which rats received DBS at 12 Hz, 60 Hz, 100 Hz, and 130 Hz as well as sham stimulation (0 μA) in a within-subjects counterbalanced design. There was a significant decrease in active lever responding, but not inactive lever responding, at each frequency of DBS compared to sham simulation (Figure 1A). Because each stimulation frequency similarly attenuated cocaine seeking, we focused on the lowest frequency tested (12 Hz) for the subsequent studies. A time course of cumulative active lever responding shows that cocaine seeking is suppressed by 12 Hz stimulation relative to sham stimulation throughout the 2-hour reinstatement sessions (Figure 1B). Electrode placements are shown in Figure 1C. Taken together, these data indicate DBS attenuates cocaine-primed reinstatement of cocaine seeking similarly across a wide variety of stimulation frequencies.

**Figure 1:**
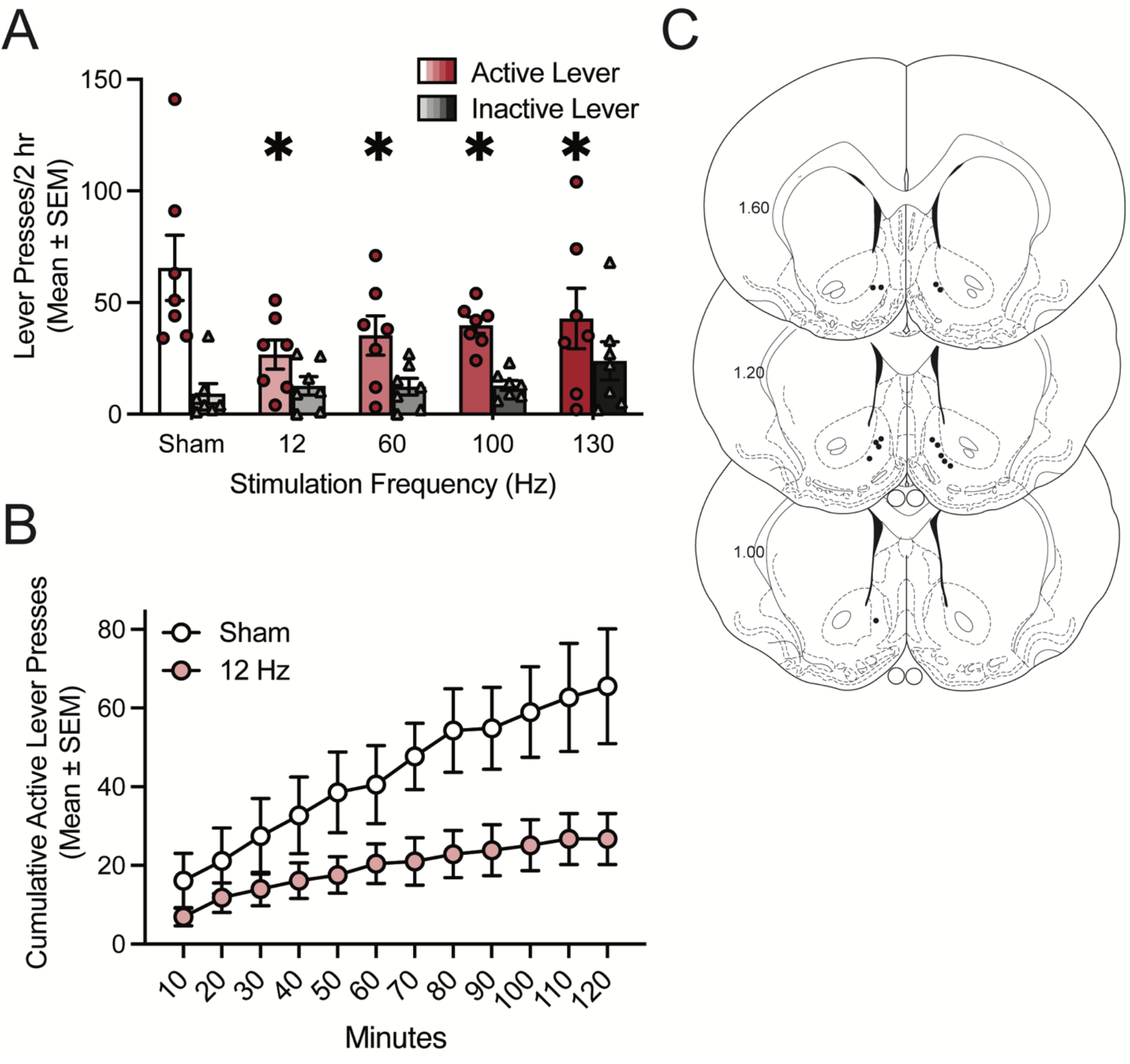
Deep brain stimulation of the nucleus accumbens attenuates cocaine seeking in male rats across a range of stimulation frequencies. A) Bars show total responding (mean ± SEM) overlaid with active lever presses (circles) and inactive lever presses (triangles) for individual rats, with each rat receiving sham and each DBS frequency in a within-subjects design. Each DBS frequency significantly attenuated active lever responding in a cocaine-primed reinstatement test (n=7). Repeated measures two-way ANOVA showed that there was no significant main effect of DBS (F_4,24_=1.66, p=0.1917), a significant main effect of lever (F_1,6_=19.82, p=0.0043), and a significant DBS x lever interaction (F_4,24_=4.12, p=0.0111). *p<0.05 vs. sham (0 µA) by Dunnett’s preplanned comparison. B) Time course of cumulative active lever presses throughout cocaine-primed reinstatement sessions where rats received sham stimulation or 12 Hz DBS. C) Electrode placements from the nucleus accumbens shell (dark circles). Values are in millimeters, relative to bregma.

### Low frequency electrical stimulation evokes long term potentiation in D1DR-MSNs and D2DR-MSNs from naive male and female rats

*Ex vivo* slice electrophysiology experiments were performed to evaluate the mechanistic effects of low frequency electrical stimulation on neuronal firing and synaptic plasticity. We first performed these experiments in naive rats to establish whether there were baseline differences by MSN subtype or sex. The effect of 12 Hz electrical stimulation on synaptic plasticity in D1DR-MSNs and D2DR-MSNs was assessed in nucleus accumbens shell Cre-dependent eYFP-positive MSNs from cocaine naive male and female D1DR-Cre and D2DR-Cre rats. The 12 Hz stimulation protocol evoked LTP in D1DR-MSNs (Figure 2A) from male (Figure 2B) and female (Figure 2C) D1DR-Cre rats as well as in D2DR-MSNs (Figure 2D) from male (Figure 2E) and female (Figure 2F) D2DR-Cre rats. Low frequency electrical stimulation also elicited specific changes in synaptic release probability (measured by paired pulse ratio, PPR) and spontaneous activity (Figure S1). Thus, low frequency electrical stimulation similarly evoked LTP across sex and MSN subtype in naive rats.

**Figure 2:**
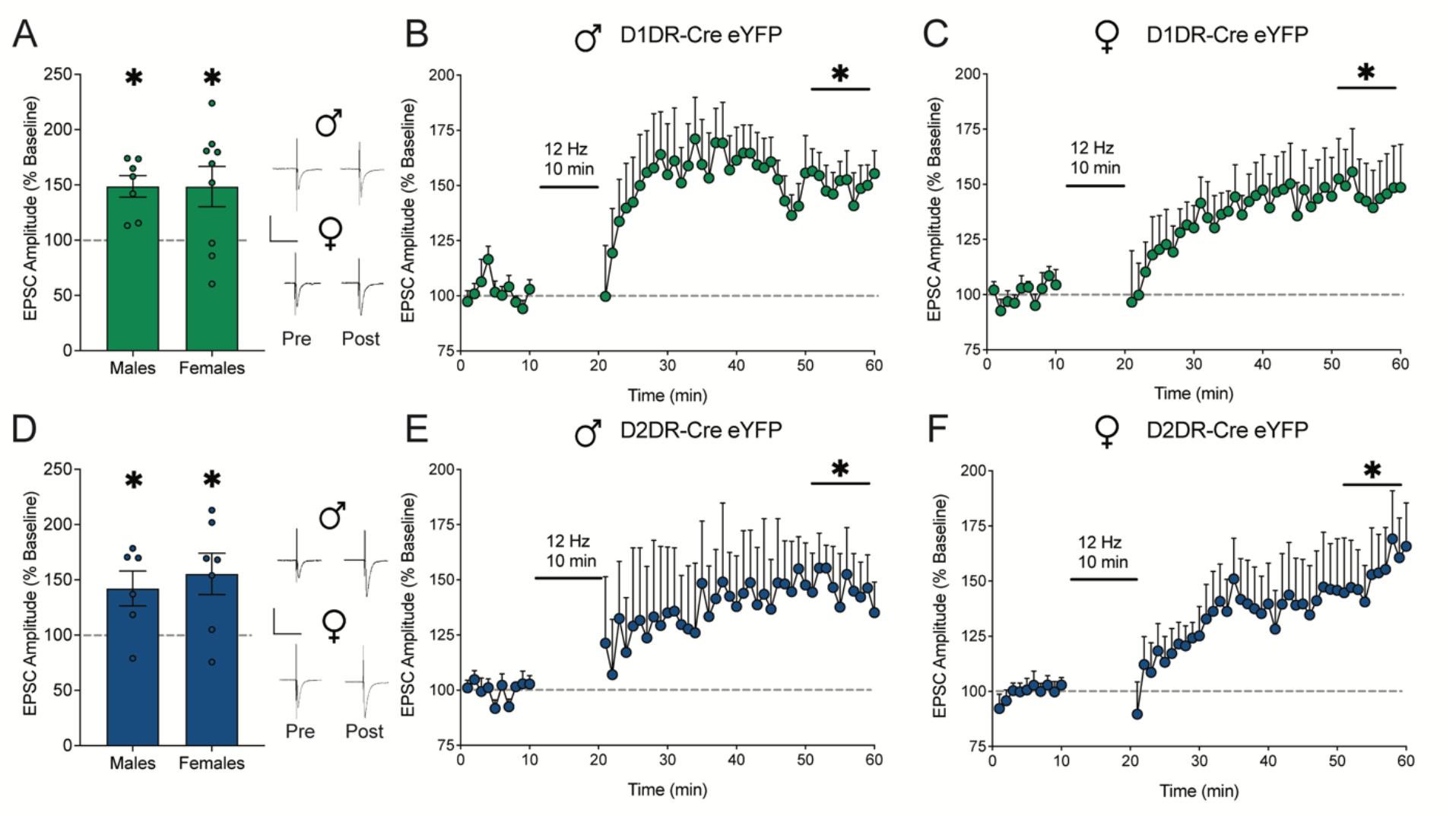
Low frequency electrical stimulation similarly elicits long term potentiation in D1DR-MSNs and D2DR-MSNs from cocaine naive male and female rats. A) EPSC amplitude averaged over the last 10 mins post-stimulation relative to 10 mins pre-stimulation baseline and full time course data for B) male (t_1,6_=4.99, p=0.0025; n=7 cells/4 rats) and C) female (t_1,8_=2.65, p=0.0291; n=9 cells/5 rats) rats expressing eYFP in D1DR-MSNs. D) EPSC amplitude averaged over the last 10 mins post-stimulation relative to 10 mins pre-stimulation baseline and full time course data for E) male (t_1,5_=2.68, p=0.044; n=6 cells/3 rats) and F) female (t_1,6_=2.95, p=0.0255; n=7 cells/4 rats) rats expressing eYFP in D2DR-MSNs. Electrical stimulation at 12 Hz evoked LTP in D1DR-MSNs and D2DR-MSNs from cocaine naive male and female rats. *p<0.05 last 10 mins post-stimulation vs. 10 mins pre-stimulation baseline by two-tailed paired t test. Insets show representative traces. Scale is 500 pA over 50 ms.

### Low frequency electrical stimulation selectively evokes long term potentiation in D2DR-MSNs from cocaine-experienced male rats

We next evaluated the effect of low frequency electrical stimulation on LTP in cocaine-experienced rats. Male (Figure 3A) and female (Figure 3B) D1DR-Cre and D2DR-Cre rats were trained to self-administer cocaine for 21 days. Lever pressing behavior was subsequently extinguished to criterion and to match the timepoint at which rats in behavioral experiments would undergo reinstatement testing. Slices from the nucleus accumbens shell were collected for electrophysiological recordings the next day.

**Figure 3:**
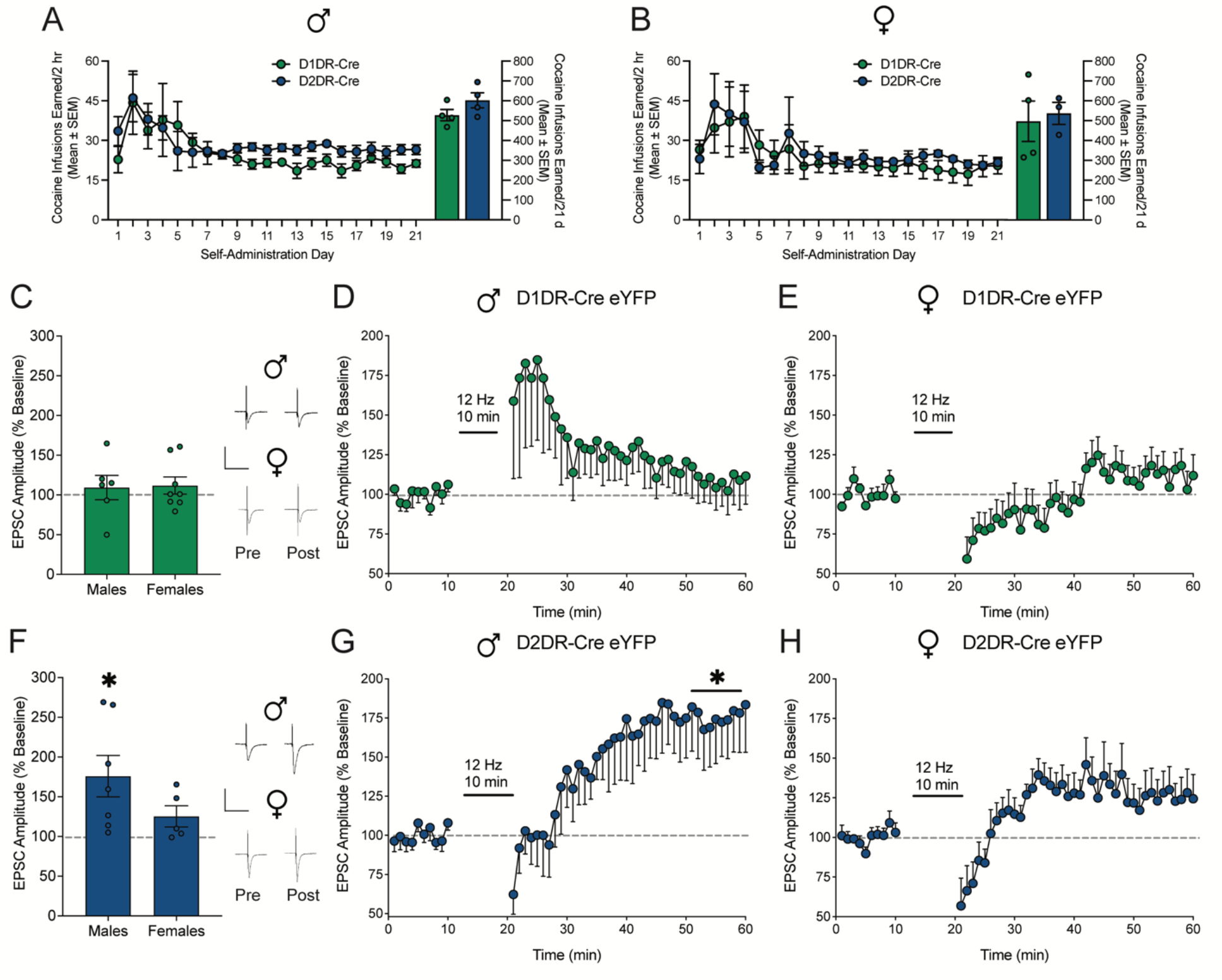
Low frequency electrical stimulation selectively elicits long term potentiation in D2DR-MSNs from cocaine-experienced male rats. Cocaine infusions earned by A) male (t_1,6_=1.574; p=0.1666) and B) female D1DR-Cre and D2DR-Cre rats did not differ (t_1,5_=0.31; p=0.77). C) EPSC amplitude averaged over the last 10 mins post-stimulation relative to 10 mins pre-stimulation baseline and full time course data for D) male (n=6 cells/4 rats) and E) female (n=8 cells/4 rats) rats expressing eYFP in D1DR-MSNs. Electrical stimulation at 12 Hz did not evoke LTP in D1DR-MSNs from cocaine-experienced male (t_1,5_=0.61, p=0.5705) and female (t_1,7_=1.09, p=0.3107) rats. F) EPSC amplitude averaged over the last 10 mins post-stimulation relative to 10 mins pre-stimulation baseline and full time course data for G) male (n=7 cells/4 rats) and H) female (n=5 cells/3 rats) rats expressing eYFP in D2DR-MSNs. Electrical stimulation at 12 Hz selectively evoked LTP in D2DR-MSNs from cocaine experienced male (t_1,6_=1.91, p=0.027), but not female rats (t_1,4_=1.90, p=0.1306). *p<0.05 last 10 mins post-stimulation vs. 10 mins pre-stimulation baseline. Insets show representative traces by two-tailed paired t test. Scale is 500 pA over 50 ms.

The effect of 12 Hz electrical stimulation on LTP was assessed in D1DR-MSNs and D2DR-MSNs labeled by Cre-dependent eYFP in D1DR-Cre and D2DR-Cre rats. Low frequency electrical stimulation failed to elicit LTP in D1DR-MSNs (Figure 3C) from male (Figure 3D) or female (Figure 3E) cocaine-experienced rats. Failure to induce LTP may reflect cocaine-mediated potentiation in D1DR-MSNs (40), which could impede the ability of electrical stimulation to further augment plasticity. Alternately, cocaine has been shown to prevent LTP in D1DR-MSNs (40), which may not be reversed under these stimulation parameters. However, in D2DR-MSNs from cocaine-experienced rats (Figure 3F), 12 Hz electrical stimulation evoked LTP in males (Figure 3G) but not females (Figure 3H). Cocaine has been shown to occlude plasticity within D2DR-MSNs (41, 42), and these data suggest 12 Hz stimulation restores LTP within this subpopulation. Differences in LTP were not concordant with alterations in PPR or spontaneous activity; only D1DR-MSNs from cocaine-experienced males and females expressed a decrease in PPR following electrical stimulation, which may suggest that DBS alters presynaptic input selectively onto D1DR-MSNs in cocaine-trained rats (Figure S2). These data indicate low frequency electrical stimulation selectively elicits LTP in D2DR-MSNs from male cocaine-experienced rats, which could be one mechanism by which 12 Hz DBS suppresses cocaine seeking.

### Low frequency optical stimulation selectively evokes long term potentiation in D2DR-MSNs from cocaine-experienced male rats

Electrical stimulation influences multiple cell types, perhaps differentially. To examine specific effects on accumbens output neurons, we used selective optogenetic low frequency stimulation of accumbens D1DR-MSNs and D2DR-MSNs. Male (Figure 4A) and female (Figure 4B) D1DR-Cre and D2DR-Cre rats were trained to self-administer cocaine for 21 days. The effect of 12 Hz optogenetic stimulation on LTP and PPR was assessed in D1DR-MSNs or D2DR-MSNs from rats which were trained to self-administer cocaine and lever responding was subsequently extinguished. D1DR-Cre and D2DR-Cre rats expressed Cre-dependent eYFP or ChR2 (with an eYFP tag) in the nucleus accumbens shell. In D1DR-MSNs (Figure 4C), 12 Hz optical stimulation did not evoke LTP in male (Figure 4D) or female (Figure 5E) cocaine-experienced D1DR-Cre rats which expressed Cre-dependent ChR2.

**Figure 4:**
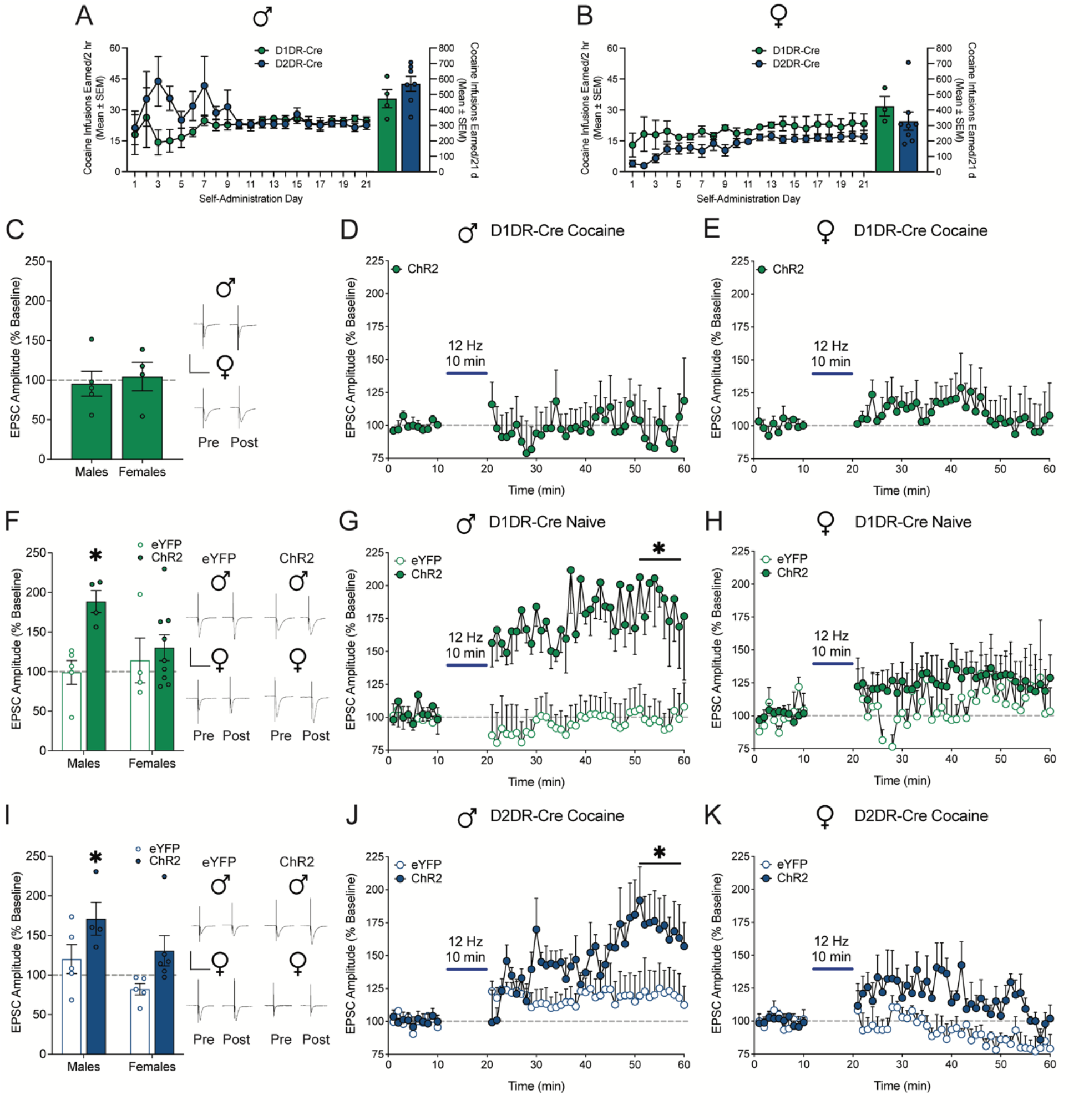
Low frequency optical stimulation selectively elicits long term potentiation in D2DR-MSNs from cocaine-experienced male rats. Cocaine infusions earned by A) male (t_1,9_=1.24; p=0.2455) and B) female D1DR-Cre and D2DR-Cre rats did not differ (t_1,9_=0.92; p=0.3857). C) EPSC amplitude averaged over the last 10 mins post-stimulation relative to 10 mins pre-stimulation baseline and full time course data for cocaine-experienced D) male (n=5 cells/4 rats) and E) female (n=4 cells/3 rats) rats expressing ChR2 in D1DR-MSNs. Optical stimulation at 12 Hz did not evoke LTP in D1DR-MSNs from cocaine-experienced male (t_1,4_=0.26, p=0.8090) and female rats (t_1,3_=0.24, p=0.8241). F) EPSC amplitude averaged over the last 10 mins post-stimulation relative to 10 mins pre-stimulation baseline and full time course data for cocaine naive G) male (eYFP n=5 cells/4 rats; ChR2 n=4 cells/3 rats) and H) female (eYFP n=4 cells/3 rats; ChR2 n=9 cells/6 rats) rats expressing eYFP or ChR2 in D1DR-MSNs. Mixed measures two-way ANOVA with stimulation (vs. baseline) and vector (eYFP vs. ChR2) as factors was used to assess the effect of 12 Hz optical stimulation on LTP. Optical stimulation at 12 Hz selectively evoked LTP in D1DR-MSNs from cocaine experienced male, but not female, rats expressing ChR2. There was no change from baseline in male or female controls that expressed eYFP alone. In male cocaine naive D1DR-Cre rats, there was a significant main effect of stimulation (F_1,7_=16.24, p=0.0050), a main effect of vector (F_1,7_=19.52, p=0.0031), and a significant stimulation x vector interaction (F_1,7_=17.02, p=0.0044). In female cocaine naive D1DR-Cre rats, there was no main effect of stimulation (F_1,11_=2.02, p=0.1833), no main effect of vector (F_1,11_=0.30, p=0.5968), and no stimulation x vector interaction (F_1,11_=0.25, p=0.6269). I) EPSC amplitude averaged over the last 10 mins post-stimulation relative to 10 mins pre-stimulation baseline and full time course data for cocaine-experienced J) male (eYFP n=5 cells/4 rats; ChR2 n=4 cells/3 rats) and K) female (eYFP n=5 cells/3 rats; ChR2 n=6 cells/4 rats) rats expressing eYFP or ChR2 in D2DR-MSNs. Optical stimulation at 12 Hz selectively evoked LTP in D2DR-MSNs from cocaine experienced male, but not female, rats expressing ChR2. There was no change from baseline in male or female controls that expressed eYFP alone. In male cocaine-experience D2DR-Cre rats, there was a main effect of stimulation (F_1,7_=9.54, p=0.0176), a main effect of vector (F_1,7_=9.52, p=0.0177), and a significant stimulation x vector interaction (F_1,7_=9.80, p=0.0209). In female cocaine-experience D2DR-Cre rats, there was no main effect of stimulation (F_1,9_=0.32, p=0.5826), a trend toward a main effect of vector (F_1,9_=4.95, p=0.0531), and a trend toward a stimulation x vector interaction (F_1,9_=4.679, p=0.0588). *p<0.05 last 10 mins post-stimulation vs. 10 mins pre-stimulation baseline by Bonferroni post-hoc analysis. Insets show representative traces. Scale is 500 pA over 50 ms.

**Figure 5:**
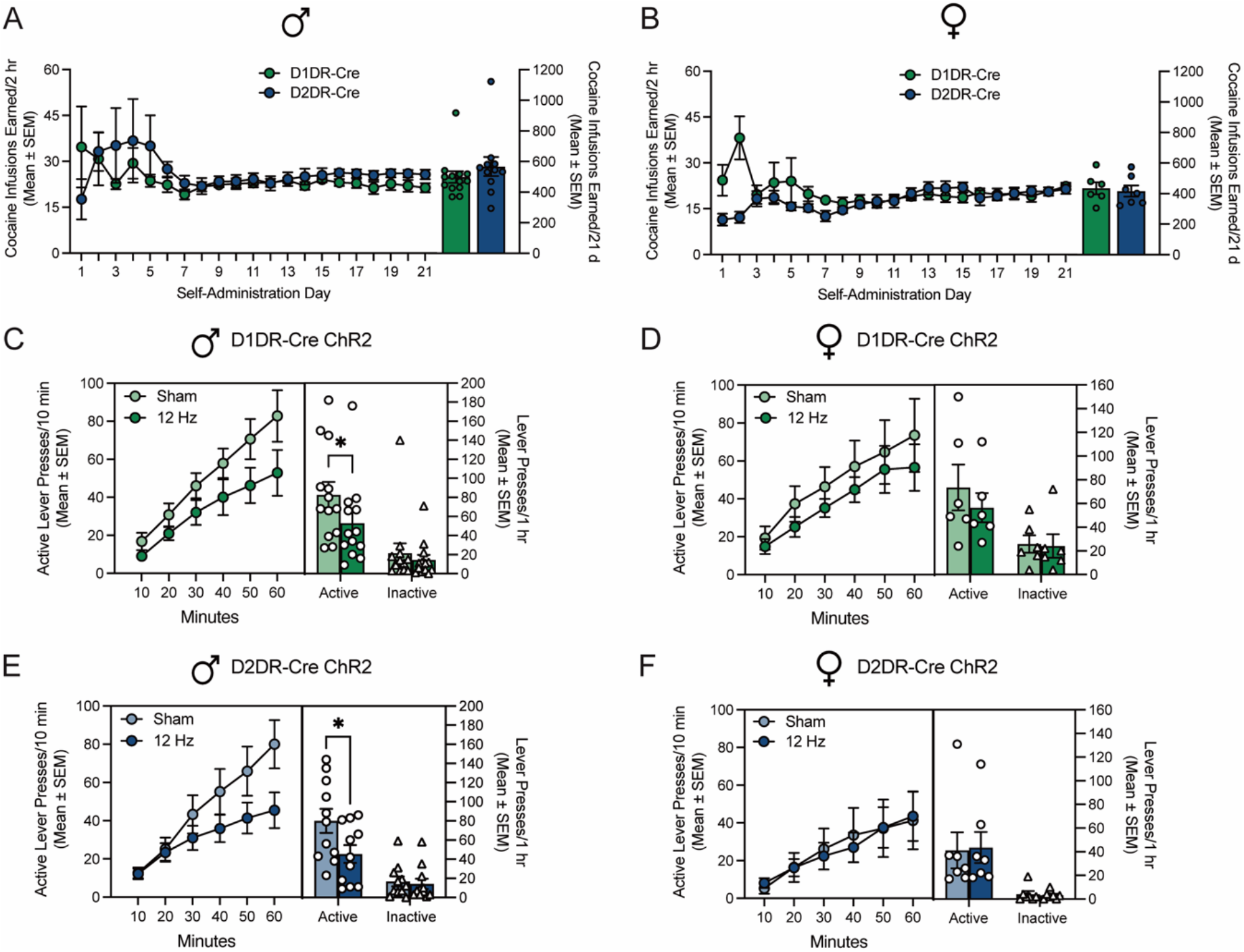
Low frequency opto-DBS in D1DR-containing and D2DR-containing neurons of the accumbens shell attenuated cocaine-primed reinstatement in male but not female rats. Cocaine infusions earned by A) male (t_1,22_=0.97; p=0.3428) and B) female D1DR-Cre and D2DR-Cre rats did not differ (t_1,11_=0.34; p=0.7371). Time courses show cumulative active lever presses during 1 hour reinstatement sessions. Bars show total responding overlaid with active lever presses (circles) and inactive lever presses (triangles) for individual rats, with each rat receiving sham and 12 Hz stimulation in a within-subjects design. C) In male rats that expressed ChR2 in D1DR-containing neurons in the nucleus accumbens shell (n=13), cocaine seeking was significantly attenuated by 12 Hz opto-DBS stimulation, relative to sham stimulation (\**p*<0.05). Two-way repeated measures ANOVA revealed a trend toward a main effect of stimulation (F_1,12_=4.03, p=0.0676), a main effect of lever (F_1,12_=23.95, p=0.0004), and no lever by stimulation interaction (F_1,12_=3.12, p=0.1029) on lever presses during the reinstatement sessions. D) In female rats that expressed ChR2 in D1DR-containing neurons in the nucleus accumbens shell (n=6), cocaine seeking did not differ when rats received sham stimulation or 12 Hz opto-DBS stimulation throughout the cocaine-primed reinstatement session. There was no main effect of stimulation (F_1,5_=0.75, p=0.4263), a main effect of lever (F_1,5_=7.13, p=0.0444), and no lever by stimulation interaction (F_1,5_=0.97, p=0.3707) on lever presses during the reinstatement sessions. E) In male rats that expressed ChR2 in D2DR-containing neurons in the nucleus accumbens shell (n=11), cocaine seeking was significantly attenuated by 12 Hz opto-DBS stimulation, relative to sham stimulation. Two-way repeated measures ANOVA revealed a trend toward a main effect of stimulation (F_1,10_=4.21, p=0.0672), a main effect of lever (F_1,10_=61.20, p<0.0001), and a trend toward a lever by stimulation interaction (F_1,10_=4.89, p=0.0515) on responses during the reinstatement sessions. F) In female rats that expressed ChR2 in D2DR-containing neurons in the nucleus accumbens shell (n=7), cocaine seeking did not differ when rats received sham stimulation or 12 Hz opto-DBS stimulation throughout the cocaine-primed reinstatement session. There was no main effect of stimulation (F_1,6_=0.031, p=0.8665), a main effect of lever (F_1,6_=7.44, p=0.0343), and no lever by stimulation interaction (F_1,6_=0.24, p=0.6407) on lever presses during the reinstatement sessions. *p<0.05 12 Hz stimulation vs. sham by Bonferroni post-hoc analysis.

To verify that this failure of optical stimulation to elicit LTP is cocaine-dependent, the effect of 12 Hz optical stimulation was assessed in cocaine naive D1DR-Cre rats which expressed Cre-dependent eYFP or ChR2. Similar to low frequency electrical stimulation, 12 Hz optical stimulation evoked LTP in male (Figure 4F, G) but not female D1DR-Cre rats expressing ChR2 (Figure 4F,H). There was no effect of optical stimulation in eYFP controls (Figure 4F-H). Thus, the inability of 12 Hz optical stimulation to elicit LTP in D1DR-MSNs is driven by a history of cocaine self-administration and not the failure of optical stimulation to evoke LTP in this population of neurons.

In D2DR-MSNs (Figure 4I), 12 Hz optical stimulation selectively evoked LTP in male (Figure 4J) but not female (Figure 4K) cocaine-experienced D2DR-Cre rats which expressed Cre-dependent ChR2, but not eYFP. Optical stimulation did not alter synaptic release probability in any group (Figure S3). These results mirror the effects of electrical stimulation and suggest that low frequency optical stimulation selectively elicits LTP in D2DR-MSNs from male cocaine-trained rats.

### Low frequency opto-DBS of accumbens D1DR- and D2DR-containing neurons attenuates cocaine seeking in male but not female rats

Lastly, the behavioral effect of low frequency opto-DBS in D1DR- and D2DR-expressing neurons to modulate cocaine seeking in male and female rats was explored. Male (Figure 5A) and female (Figure 5B) rats expressing Cre-dependent ChR2 in D1DR-containing or D2DR- containing neurons self-administered cocaine for 21 days, after which lever responding was extinguished. Cre Rats then received 12 Hz opto-DBS and sham stimulation throughout 1-hour cocaine-primed reinstatement tests in a within-subjects counterbalanced fashion. In rats that expressed ChR2 in nucleus accumbens shell D1DR-containing neurons, low frequency opto-DBS suppressed cocaine priming-induced reinstatement of drug seeking in male rats (Figure 5C) but not female rats (Figure 5D). Similarly, in rats that expressed ChR2 in nucleus accumbens shell D2DR-containing neurons, low frequency opto-DBS suppressed cocaine priming-induced reinstatement of drug seeking in male rats (Figure 5E) but not female rats (Figure 5F). Collectively, these data indicate that low frequency optogenetic stimulation of D1DR- and D2DR-containing neurons attenuates cocaine priming-induced reinstatement in male rats, similar to that of electrical DBS stimulation (Figure 1, (23)). However, low frequency optogenetic stimulation of D1DR-containing or D2DR-containing does not alter cocaine seeking in female rats, similar to prior work with high frequency optogenetic stimulation (28).

## Discussion

The present findings demonstrated that low frequency DBS of the nucleus accumbens shell attenuated cocaine seeking in male rats. While 12 Hz electrical stimulation elicited LTP in both D1DR-MSN and D2DR-MSN subtypes from naive male and female rats, a history of cocaine self-administration and extinction training occluded this plasticity such that low frequency stimulation only evoked LTP in D2DR-MSNs from male cocaine-experienced rats.

Selective optical stimulation of a subpopulation of cells replicated this effect such that LTP was only produced in D2DR-MSNs from male cocaine-trained rats, although low frequency optical stimulation in D1DR-MSNs from cocaine naive male rats also was able to elicit LTP. Low frequency opto-DBS of accumbens D1DR- and D2DR-containing accumbens neurons similarly attenuated cocaine-primed reinstatement of cocaine seeking in male rats despite these subtype-specific differences in the induction of long-term synaptic plasticity. Low frequency opto-DBS of either subpopulation of neurons failed to influence cocaine seeking in female rats similar to our recent work using high-frequency opto-DBS (28), which indicates that there may be important sex differences in the efficacy of accumbens DBS. The present results indicate that low frequency DBS may suppress cocaine seeking in males via actions on both D1DR-MSNs and D2DR-MSNs, although the cellular mechanisms by which stimulation influences behavior appear to be distinct.

We expected that selective 12 Hz stimulation of D2DR-containing neurons would attenuate cocaine seeking given that i) high frequency opto-DBS of D2DR-containing neurons but not D1DR-containing neurons suppressed cocaine reinstatement (28) and that ii) 12 Hz stimulation evoked LTP in D2DR-MSNs, but not cocaine-exposed D1DR-MSNs. Surprisingly, cocaine seeking was similarly attenuated by 12 Hz opto-DBS of both D1DR- and D2DR-containing neurons. Mimicking DBS with low frequency optogenetic stimulation of prefrontal cortex inputs onto accumbens D1DR-MSNs has previously been shown to block cocaine sensitization and produce long term depression (LTD) (41). Optogenetic stimulation replicated the behavioral effects of high frequency DBS administered alone and co-administration of 12 Hz electrical DBS with a D1DR antagonist (29). In terms of outputs of accumbens MSNs, cocaine blocked LTP in D1DR-MSNs which projected to the ventral pallidum (41), similar to the present study. A stimulation protocol which induced LTD in naive D1DR-MSNS rescued cocaine-occluded LTP in D1DR-MSN outputs and also blocked cocaine sensitization (41). Future work might focus on varying stimulation parameters to reverse cocaine-mediated occlusion of LTP in D1DR-MSNs and exploring the effect of such DBS protocols on cocaine seeking. Together with the present results, these studies support the idea that activation of the D1DR-containing subpopulation of MSNs can attenuate cocaine-related behaviors under certain conditions, including specific stimulation parameters and within specific pathways (27). It also is important to establish a causal link between DBS-mediated LTP rescue in D2DR-MSNs and attenuation of cocaine-seeking. These findings underscore the importance of understanding the subtype-specific effects of stimulation parameters to optimize the therapeutic effect of DBS for substance use disorders.

Electrical DBS delivered at high frequencies induces a wide range of neurophysiological changes, including activation (43, 44), inhibition locally or via antidromic stimulation (45-48), and/or circuit-wide action on fibers of passage (48-52). Here we show that low frequency (12 Hz) electrical stimulation evoked LTP in both D1DR-MSNs and D2DR-MSNs in naive rats. In rats that had self-administered cocaine and subsequently undergone extinction of lever pressing to criterion, LTP was elicited only in D2DR-MSNs from male rats. It is notable that this experimental design, used to replicate the neuronal state of rats at the time of cocaine reinstatement sessions, differs from prior work using acute or repeated non-contingent cocaine delivery paradigms (e.g., (29, 40, 41)). A few studies have examined MSN subtype-specific aspects of accumbens synaptic plasticity after cocaine self-administration and withdrawal (53, 54); however, extinction learning following cocaine self-administration alters neuronal plasticity within accumbens MSNs (55-57) and may alter the effect of DBS on cocaine seeking. A caveat in drawing comparisons across our electrophysiology and behavior results is that rats evaluated for cocaine seeking during intra-accumbens DBS or opto-DBS also receive an acute injection of cocaine to reinstate lever pressing. We have not yet established how acute re-exposure to cocaine might influence stimulation-evoked LTP. Mimicking DBS with optogenetics allows isolation of the effect of stimulation directly on nucleus accumbens MSNs and excludes the range of other effects of electrical DBS that are likely to impinge upon neuronal firing. In this context, any effects on presynaptic release are mediated specifically by retrograde activation or inhibition from the postsynaptic MSN. It would be interesting to distinguish the effect of retrograde signaling from other presynaptic mechanisms of electrical DBS. As one example, presynaptic release probability is reduced following both 12 Hz electrical and optical stimulation in D1DR-MSNs (Figures S1 and S2) and retrograde signaling from this subpopulation may contribute to the mechanisms by which opto-DBS of D1DR-containing neurons suppresses cocaine seeking.

The absence of an effect of low frequency opto-DBS on cocaine seeking in female rats replicates our prior work with high frequency opto-DBS (28). It is important to note that the ability of electrical DBS to alter cocaine seeking in female rats is yet unknown. In cocaine-experienced rats, 12 Hz optical stimulation was able to elicit LTP in D2DR-MSNs from cocaine-experienced male, but not female rats. The present study has not causally linked DBS-evoked LTP in MSN subtypes to cocaine seeking; however, sex differences in synaptic plasticity may underlie these behavioral differences. Optical stimulation did not evoke LTP in D1DR-MSNs from cocaine-experienced male or female rats, but only male rats showed attenuated cocaine seeking during opto-DBS of D1DR-MSNs. In naive rats, 12 Hz optical stimulation was only able to elicit LTP in D1DR-MSNs from male rats, despite the fact that LTP was observed in both sexes following electrical stimulation. This could point to a distinction between the ways in which electrical DBS, which includes indirect presynaptic effects not replicated by direct optical stimulation of accumbens MSNs, produces similar effects on LTP between males and females. Sex differences in excitability within key accumbens afferent (e.g., prefrontal cortex, ventral hippocampus) and/or efferent (e.g., ventral pallidum, ventral tegmental area) pathways may also govern the effect of accumbens stimulation. For example, testosterone regulates excitability in the ventral hippocampus to nucleus accumbens pathway via local androgen receptors, and female mice exhibit inherent increased excitability within this circuit (58). Inherent hyperexcitability in females may reduce the ability of DBS-like stimulation to evoke changes in plasticity and suppress cocaine seeking. Cocaine self-administration and extinction/reinstatement were performed identically across male and female rats in the present study; consideration of additional factors like gonadal hormones and estrous cycle may identify parameters in which DBS and opto-DBS effectively suppress cocaine seeking in females (59-61).

Our work contributes to a growing literature which supports the use of DBS for substance use disorders (2-5). The present study indicates that lower DBS frequencies are similarly efficacious in the suppression of craving, although the mechanisms by which high vs. low frequency stimulation alter behavior may be distinct. Refinement of DBS approaches by altering frequencies of stimulation or targeting distinct mechanisms (e.g., plasticity within neuronal subtypes) may improve treatment outcomes. Our study also highlights unexpected and dramatic sex differences in the ability of accumbens DBS to attenuate drug craving, which is an important consideration when translating results such as these to the clinic.

## Competing Interest

The authors declare no competing interests.

## Author Contributions

SESJ and RCP conceptualized the study, SESJ, MTR, and RCP designed experiments; SESJ, MTR, PJH, MCK, AST, SM, and SJW performed experiments; SESJ, MTR, and RCP analyzed data; SESJ wrote the first draft; SESJ, MTR, and RCP wrote the manuscript; all authors reviewed and approved the manuscript.

## Acknowledgements

This work was supported by grants from the National Institutes of Health R01 DA015214 (RCP), T32 DA028874 (SESJ) and F32 DA052993 (MTR) as well as research funds from the Rutgers Robert Wood Johnson School of Medicine.

The authors thank Mateo Sarmiento, Riley Merkel, Tyler Sacko, Dominick Gangemi, Harsh Rohilla, Ayanna Coleman, Disha Shah, Izabella Mus, Stefanie Devizio, and Maya Abdelaziz for their technical contributions and maintenance of transgenic colonies and Zhiping Pang for advice and technical assistance.

## Notes

### Competing Interest Statement

The authors have declared no competing interest.

