## Supplemental Methods and Figures for "Low frequency optogenetic deep brain stimulation of nucleus accumbens dopamine D1 or D2 receptor-containing neurons attenuates cocaine seeking selectively in male rats in part by reversing synaptic plasticity deficits"

*Genotyping:* Long Evans transgenic D1DR-Cre or D2DR-Cre rats were bred in house and Cre expression in D1DR- or D2DR-containing neurons was confirmed by genotyping with the following primers (5' to 3'): D1DR forward - CTCCTGATGGAACCCTACCA; D2DR forward - TCAGGGAACCCTCTTTGAGA; Cre reverse - CACAGTCAGCAGGTTGGAGA (Sigma Aldrich, St. Louis, MO).

*Drugs*: Cocaine hydrochloride was obtained from the National Institute on Drug Abuse (Rockville, MD) and dissolved in bacteriostatic 0.9% saline.

*Surgery:* Prior to surgery, rats were anesthetized with 80 mg/kg ketamine and 12 mg/kg xylazine. An indwelling silastic catheter (SAI Infusion Technologies, Libertyville, Il) was inserted into the jugular vein and sutured in place as previously described [33,34]. Following catheter insertion, rats were mounted in a stereotaxic apparatus (Kopf Instruments, CA) for implantation of electrodes or viral vector infusion with fiber optic implantation. Bipolar stainless steel electrodes (P1 Technologies, Roanoke, VA) were cut to 7.3 mm and implanted in the nucleus accumbens shell at 1.0 mm A/P and ±3.0 mm M/L on a 17° angle, relative to bregma (Paxinos and Watson, 1997). Viral infusions and implantation of fiber optic targeting the nucleus accumbens shell were performed using the coordinates +1.0 mm A/P, ±3.0 mm M/L, -7.3 mm D/V on a 17° angle, relative to bregma (Paxinos and Watson, 1997). Bilateral infusions (1 μl/side) of a Cre-dependent adeno-associated viral vector (AAV) expressing eYFP (AAV5-EF1a-DIO-eYFP-WPRE-hGH; Addgene) or a Cre-dependent AAV expressing ChR2 and eYFP (AAV5-EF1a-DIO-hChR2(H134R)-eYFP-WPRE-hGH; Addgene) were delivered via Hamilton syringes (Reno, NV) into the nucleus accumbens shell. Viral vector delivery was immediately followed by placement of 200 μm fiber optic (Thor Labs, Newton, NJ) attached to stainless steel ferrules (Fiber Instrument Sales, Oriskany, NY) and cut to length to terminate just above the nucleus accumbens shell. Electrodes or ferrules were cemented in place by affixing dental acrylic to three stainless steel screws fastened to the skull. Rats recovered for at least seven days; catheters were flushed daily (Timentin, 0.93 mg/ml, in heparinized saline) during recovery and after each daily behavioral session.

*Ex vivo slice preparation:* Rats were perfused with ice-cold cutting solution containing (in mM): 92 N-methyl-d-glucamine (NMDG), 2.5 KCl, 1.2 $NaH_2PO_4$, 30 $NaHCO_3$, 20 HEPES, 25 glucose, 5 sodium ascorbate, 2 thiourea, 3 sodium pyruvate, 10 $MgSO_4$, and 0.5 $CaCl_2$, saturated with carbogen (95% O2/5% CO2), pH adjusted to 7.4 with HCl. Rats were then decapitated and brains removed. Acute coronal slices of the nucleus accumbens (250 µm thick) were obtained using a VT1000S vibratome (Leica, Weltzar, Germany) in 4°C cutting solution, then placed in a holding chamber of the same cutting solution, and incubated at 37°C for 10-12 min. Slices were then transferred to a beaker of room temperature holding ACSF containing (in mM): 86 NaCl, 2.5 KCl, 1.2 $NaH_2PO_4$, 35 $NaHCO_3$, 20 HEPES, 25 glucose, 5 sodium ascorbate, 2 thiourea, 3 sodium pyruvate, 1 $MgCl_2$, and 2 $CaCl_2$, saturated with carbogen, pH 7.3-7.4, osmolarity 305-315 mOsm. Slices were allowed to recover for at least 45 min before performing recordings.

*Ex vivo electrophysiology:* Slices were placed on a Nikon Eclipse FN1 upright microscope equipped for Differential Interference Contrast (DIC) infrared optics. The nucleus accumbens was identified using a 5X objective and individual neurons were magnified with a 40X water immersion lens. The recording chamber was continuously perfused with oxygenated recording ACSF containing (in mM): NaCl 119, KCl 2.5, $NaHCO_3$ 26, $NaH_2PO_4$ 1.2, glucose 12.5, HEPES 5, $MgSO_4$ 1, $CaCl_2$ 2, pH 7.3-7.4, osmolarity 305-315 mOsm. The recording solution was heated to 32 ± 1°C using an automatic temperature controller (Warner Instruments, Holliston MA). Picrotoxin (100 µM; dissolved in DMSO) was included in the recording solution to inhibit $GABA_A$ receptor-mediated currents. Recording pipettes were pulled from borosilicate glass capillaries (World Precision Instruments, Sarasota, FL) to a resistance of 4.0-5.3 MΩ when filled with intracellular solution.The intracellular solution contained the following (in mM) potassium gluconate 145, KCl 2.5, NaCl 2.5, BAPTA 0.1, HEPES 10, L-glutathione 1.0, sodium phosphocreatine 7.5, Mg-ATP 2.0, and Tris-GTP 0.25, pH 7.2-7.3 with KOH, osmolarity 285-295 mOsm.

*Verification of AAV expression and fiber optic or electrode placement.* After the completion of electrical DBS experiments, brains were fixed in 10% formalin. Coronal sections (100 µm) were made on a vibratome (Technical Products International; St. Louis, MO) and electrode track marks were visualized under a light microscope. After the completion of opto-DBS experiments, rats were given an overdose of pentobarbital (100 mg/kg) and perfused intracardially with 0.9% saline followed by 4% paraformaldehyde. The brains were removed and coronal sections (40 µm) were collected for visualization on a confocal microscope (Leica Biosystems, Buffalo Grove, IL). Alternately, rats were decapitated and viral placement was visualized using NIGHTSEA™ DFP™ Dual Fluorescent Protein Excitation Flashlight (Electron Microscopy Services, Hatfield, PA). Rats lacking Cre-dependent eYFP fluorescence or with fluorescence and/or fiber optic or electrode placement outside of the nucleus accumbens shell were excluded from subsequent data analysis.

**Supplemental Figures and Figure Legends:**

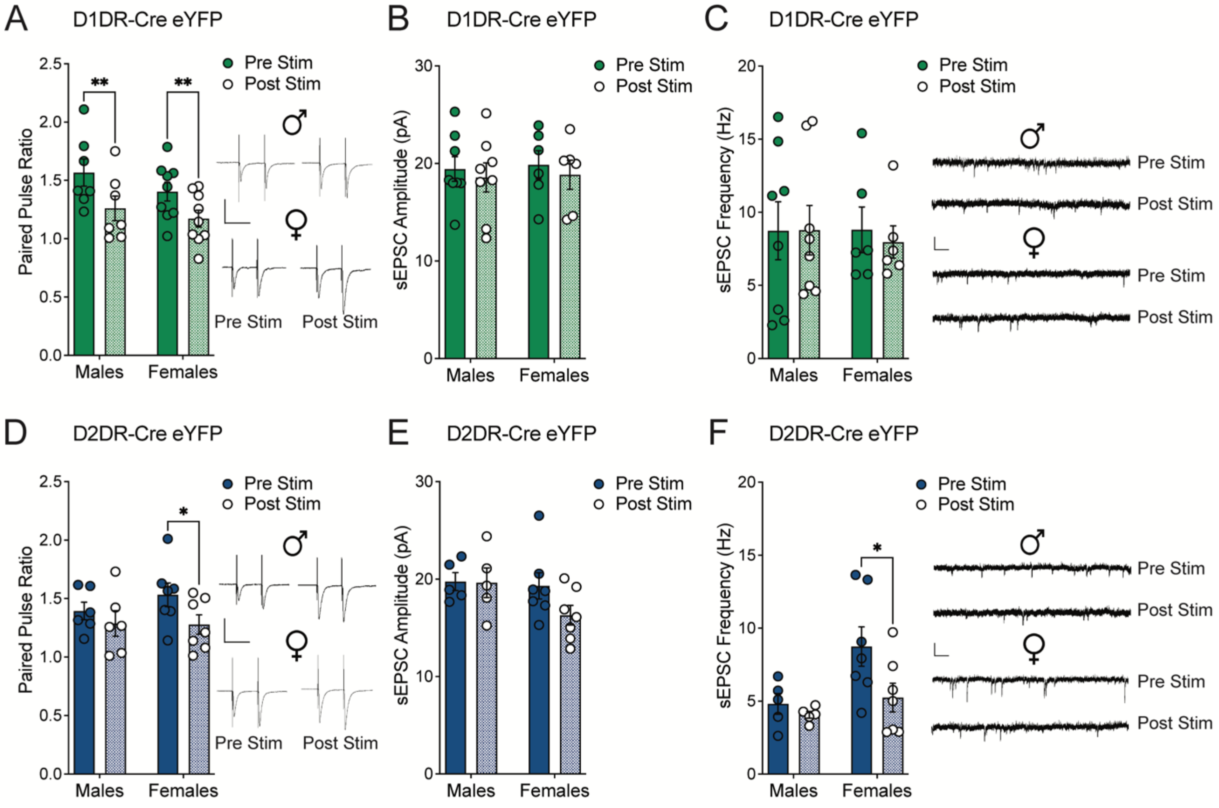


**Supplemental Figure 1:** *Low frequency electrical stimulation alters synaptic release probability and spontaneous activity in a sex- and MSN subtype-dependent manner.* A) Paired pulse ratio (PPR) was significantly lower after 12 Hz electrical stimulation in D1DR-MSNs from cocaine naive male ($t_{1,6}$=3.53, p=0.0124; n=7 cells/4 rats) female rats ($t_{1,8}$=4.54, p=0.0019; n=9 cells/5 rats). B) Spontaneous EPSC amplitude and C) frequency were not altered by electrical stimulation in D1DR-MSNs from male (amplitude: $t_{1,7}$=0.46, p=0.6567; frequency: $t_{1,7}$=0.034, p=0.9739; n=8 cells/5 rats) or female rats (amplitude: $t_{1,5}$=0.48, p=0.6507; frequency: $t_{1,5}$=0.39, p=0.7148; n=6 cells/3 rats). D) PPR was significantly lower after 12 Hz electrical stimulation in D2DR-MSNs from cocaine naive female ($t_{1,6}$=2.87, p=0.0285; n=7 cells/4 rats), but not male rats ($t_{1,5}$=1.83, p=0.1263; n=6 cells/3 rats). E) Spontaneous EPSC amplitude was not altered by electrical stimulation in D2DR-MSNs from male ($t_{1,4}$=0.064, p=0.9521; n=5 cells/3 rats) or female rats ($t_{1,6}$=2.06, p=0.0851; n=7 cells/6 rats). F) Spontaneous EPSC frequency was significantly lower after 12 Hz electrical stimulation in D2DR-MSNs from cocaine naive female ($t_{1,6}$=2.75, p=0.0335; n=7 cells/6 rats), but not male rats ($t_{1,4}$=0.98, p=0.3808; n=5 cells/3 rats). This effect may be driven by a sex difference in spontaneous EPSC frequency pre-stimulation such that female rats had higher sEPSC frequency in D2DR-MSNs than males at baseline. *$p<0.05$ post-stimulation vs. pre-stimulation by two-tailed paired t test. Insets show representative traces. Scale is 500 pA over 50 ms for PPR (A, D); 40 pA over 200 ms for spontaneous activity (B, C, E, F).

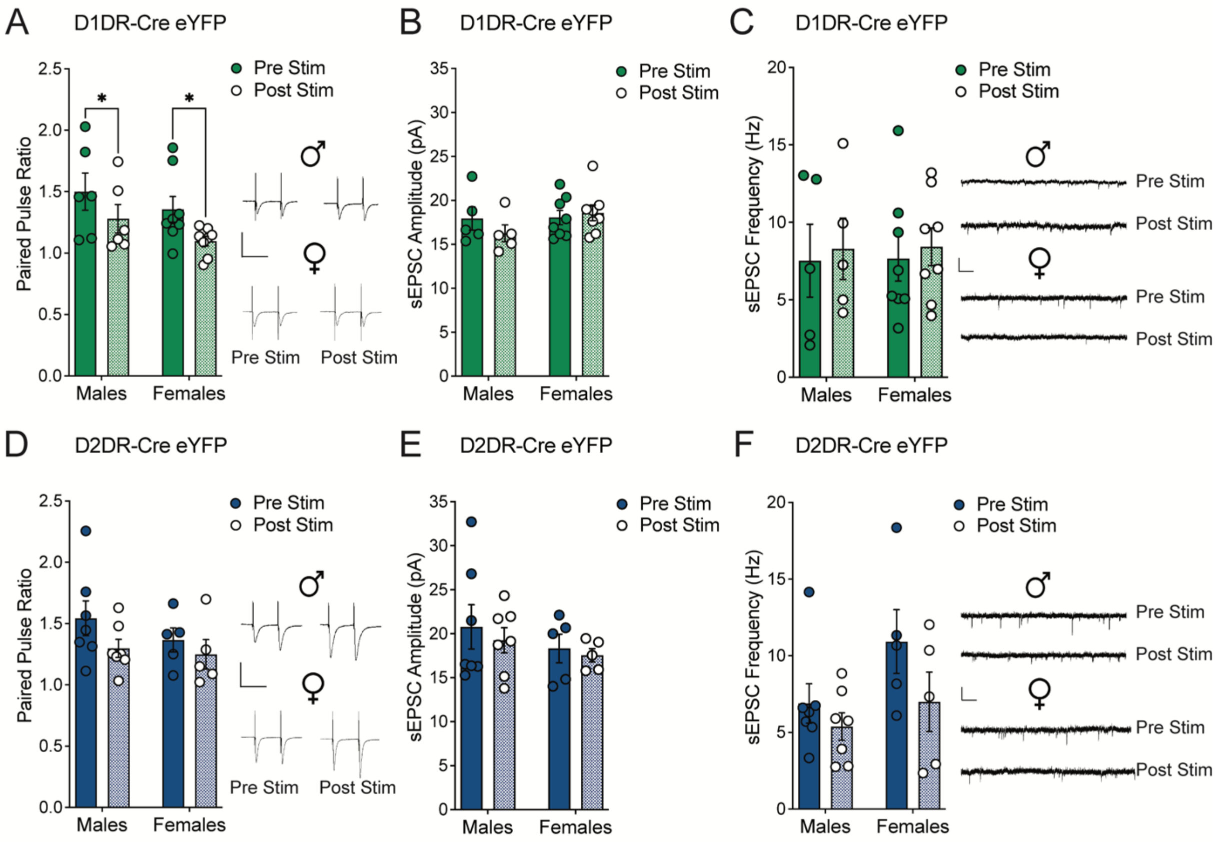


**Supplemental Figure 2:** *Effect of low frequency electrical stimulation on presynaptic release probability and spontaneous activity in D1DR-MSNS and D2DR-MSNs from cocaine-experienced male and female rats.* A) Paired pulse ratio (PPR) was significantly lower after 12 Hz electrical stimulation in cocaine-experienced male D1DR-MSNs ($t_{1,5}=3.35$, $p=0.0203$; n=6 cells/4 rats) and female D1DR-MSNs ($t_{1,6}=2.60$, $p=0.0355$; n=8 cells/4 rats). B) Spontaneous EPSC amplitude and C) frequency were not altered by electrical stimulation in D1DR-MSNs from male (amplitude: $t_{1,4}=1.30$, $p=0.2646$; frequency: $t_{1,4}=0.27$, $p=0.8002$; n=5 cells/4 rats) or female cocaine-experienced rats (amplitude: $t_{1,7}=0.67$, $p=0.5239$; frequency: $t_{1,7}=0.73$, $p=0.4909$; n=8 cells/4 rats). D) PPR was not altered by 12 Hz electrical stimulation in D2DR-MSNs from cocaine-experienced male ($t_{1,6}=1.78$, $p=0.1253$; n=7 cells/4 rats) and female rats

($t_{1,4}$=1.50, p=0.2072; n=5 cells/3 rats). E) Spontaneous EPSC amplitude and F) frequency were not altered by electrical stimulation in D2DR-MSNs from male (amplitude: $t_{1,6}$=0.59, p=0.5750; frequency: $t_{1,6}$=1.76, p=0.1290; n=7 cells/4 rats) or female cocaine-experienced rats (amplitude: $t_{1,4}$=0.63, p=0.6922; frequency: $t_{1,4}$=1.59, p=0.1865; n=5 cells/3 rats). *p<0.05 post-stimulation vs. pre-stimulation by two-tailed paired t test. Insets show representative traces. Scale is 500 pA over 50 ms for PPR (A, D); 40 pA over 200 ms for spontaneous activity (B, C, E, F).

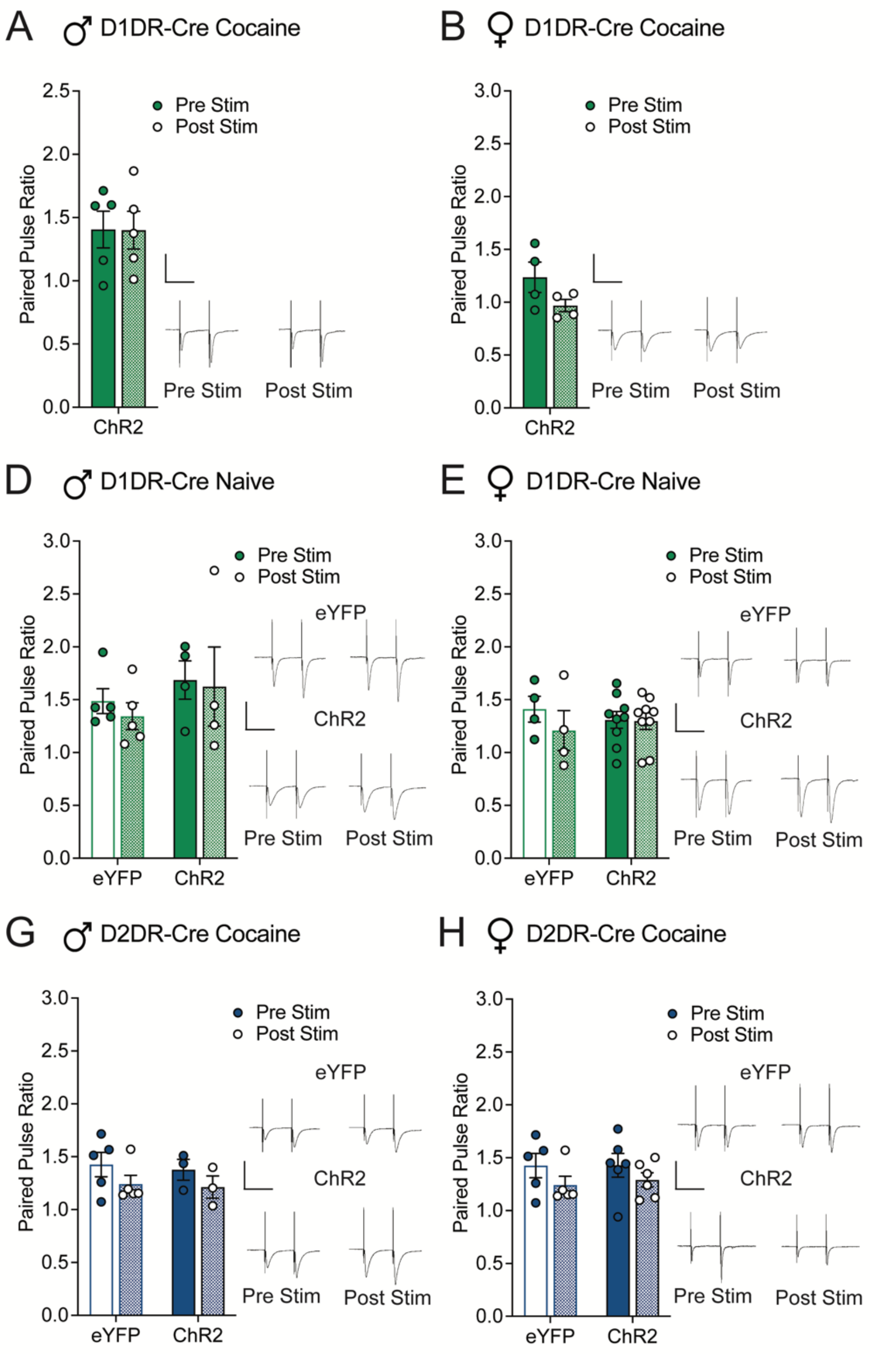


**Supplemental Figure 3:** *Low frequency optical stimulation did not alter presynaptic release probability in D1DR-MSNs and D2DR-MSNs from cocaine-experienced rats.* Paired pulse ratio (PPR) was not altered by 12 Hz optical stimulation in cocaine-experienced A) male ($t_{1,4}$=0.050, p=0.9622; n=5 cells/4 rats) and B) female ($t_{1,3}$=2.915, p=0.0617; n=4 cells/3 rats) rats expressing ChR2 in D1DR-MSNs. PPR was not altered by 12 Hz optical stimulation in cocaine naive C) male (eYFP n=5 cells/4 rats; ChR2 n=4 cells/3 rats) and D) female (eYFP n=4 cells/3 rats; ChR2 n=9 cells/6 rats) rats expressing eYFP or ChR2 in D1DR-MSNs. In cocaine naive D1DR-Cre males, two-way mixed measures ANOVA revealed that there was a trend toward a main effect of stimulation ($F_{1,7}$=0.50, p=0.5015), no main effect of vector ($F_{1,7}$=0.88, p=0.3787), and no stimulation x vector interaction ($F_{1,7}$=0.077, p=0.7895). In cocaine naive D1DR-Cre females, two-way mixed measures ANOVA revealed that there was no main effect of stimulation ($F_{1,11}$=1.60, p=0.2320), no main effect of vector ($F_{1,11}$=0.0024, p=0.9614), and no stimulation x vector interaction ($F_{1,11}$=1.26, p=0.2849). PPR was not altered by 12 Hz optical stimulation in cocaine-experienced E) male (eYFP n=5 cells/4 rats; ChR2 n=3 cells/2 rats) and F) female (eYFP n=5 cells/3 rats; ChR2 n=6 cells/4 rats) rats expressing eYFP or ChR2 in D2DR-MSNs. In cocaine-experienced D2DR-Cre males, two-way mixed measures ANOVA revealed that there was a main effect of stimulation ($F_{1,6}$=9.22, p=0.0229), no main effect of vector ($F_{1,6}$=0.075, p=0.7928), and no stimulation x vector interaction ($F_{1,6}$=0.034, p=0.8601). In cocaine-experienced D2DR-Cre females, two-way mixed measures ANOVA revealed that there was a main effect of stimulation ($F_{1,9}$=6.99, p=0.0267), no main effect of vector ($F_{1,9}$=0.050, p=0.8375), and a trend toward a stimulation x vector interaction ($F_{1,9}$=2.915, p=0.0617). Bonferroni post-hoc analyses showed that there was no significant effect of 12 Hz optical stimulation on PPR in any group. Insets show representative traces. Scale is 500 pA over 50 ms.
